# Genomic contingencies and beak shape variation in a hybrid species

**DOI:** 10.1101/107490

**Authors:** Anna Runemark, Laura Piñeiro Fernández, Fabrice Eroukhmanoff, GlennPeter Sætre

**Affiliations:** Centre for Ecological and Evolutionary Synthesis, Department of Biosciences, University of Oslo; Institut für Systematische und Evolutionäre Botanik, University of Zürich

**Keywords:** adaptation, beak shape, climate, diet, genetic constraints, Passer italiae

## Abstract

Hybridization is increasingly recognized as a potent evolutionary force. Though additive genetic variation and novel combinations of parental genes theoretically increase the potential for hybrid species to adapt, few empirical studies have investigated the adaptive potential within a hybrid species. Here, we investigate factors promoting phenotypic divergence using genomically diverged island populations of the homoploid hybrid Italian sparrow *Passer italiae* from Crete, Corsica, and Sicily. We address whether genomic contingencies, adaptation to climate or diet best explain divergence in beak morphology. Populations vary significantly in beak morphology, both between and within islands of origin. Temperature seasonality best explains population divergence in beak size. Interestingly, beak shape along all significant dimensions of variation was best explained by annual precipitation, genomic composition and their interaction, suggesting a role for contingencies. Moreover, beak shape similarity to a parent species correlates with proportion of the genome inherited from that species, consistent with the presence of contingencies. In conclusion, adaptation to local conditions and genomic contingencies arising from putatively independent hybridization events jointly explain beak morphology in the Italian sparrow. Hence, hybridization may induce contingencies and restrict evolution in certain directions dependent on the genetic background.

---

Data archival location: Dryad

## Introduction

Adaptation to divergent ecological niches is a major factor in population divergence and speciation (Schluter 2000; Grant and Grant 2008; Schluter 2009). Adaptation in key traits where novel morphologies can allow for the invasion of new niches (Dumont et al. 2012), are of particular interest since divergence in these can drive speciation (Hunter 1998). Key traits can also enable co-existence with closely related species (Miraldo and Hanski 2014) and hence spur adaptive radiations (Schluter 2000), and can generate specious groups, such as birds (Jarvis et al. 2014). The beak is such a key trait, since beak shape adaptations have significantly contributed to the niche diversity in birds (Mallarino et al. 2012). Variation in beak size- and shape is important both for feeding efficiency (Benkman 2002; 2016) and thermoregulation (Symonds and Tattersall 2010). It is also affects song (Derryberry et al. 2012), and can hence be a target of sexual selection (Huber and Podos 2006). A classical example of beak morphology adaptation is the radiation of Darwin’s finches on the Galapagos Islands, where divergent selection between groups of birds with different dietary preferences have caused a dramatic beak shape diversity (Grant and Grant 2006). Interestingly, hybridization can also generate new beak shapes that allow more efficient use of specific dietary resources (Grant and Grant 1996; Lamichhaney et al. 2015; 2016).

Hybridization is increasingly recognized as an important source of novel genetic variation (Mallet 2005; 2007; Abbott et al. 2013). It can spur novel adaptations by increasing genomic diversity, and through changing the constraints on the direction of evolution. Hybrids are expected to have more additive genetic variation than the parental species’ genomes, and this increase is highest when the parent species are fixed for different alleles at each locus (Bailey et al. 2013; Seehausen 2013; Eroukhmanoff et al. 2013b). Furthermore, the mosaic genome from the combination of the two parental genome complexes (Rieseberg 2003) can give rise either to phenotypes that are intermediate or mosaic versions of the parents, or transgressive phenotypes, which are beyond the range of the parental species (Rieseberg et al. 1999; Dittrich-Reed and Fitzpatrick 2012). The increase in additive genetic variation and the novel combinations of parental genes may increase the potential for hybrids to adapt (Rieseberg 2003; Eroukhmanoff et al. 2013b). Interestingly, different hybrid populations can attain strongly divergent genomic composition (Runemark et al. n.d.). However, hybrid species can also be subjected to constraints or contingencies resulting from mosaic patterns of parental inheritance or conditions during initial hybridization and genome stabilization (Eroukhmanoff et al. 2013b). Moreover, depending on the type of selection acting on the parent species’ phenotypes, hybrid morphology is expected to be more or less restricted. For traits under stabilizing selection in parents, hybrids are expected to be free to evolve towards a variety of different potential fitness optima, even those extending beyond those of the parents (Bailey et al. 2013). However, when directional selection has contributed to parent species differences, hybrid phenotypes are predicted to be intermediate of the parent taxa, and restricted to evolve along the axis of divergence between them (Bailey et al. 2013). This could facilitate convergence towards parental phenotypes (Bailey et al. 2013). In the latter situation, populations of hybrid species could be restricted to trait values reflecting the relative proportion of the genome inherited from the parent species. Hence, hybrid populations differing in genomic composition could either be divergent due to genomic contingencies or to adaptation in response to local selection pressures. Tests for presence of such genomic contingencies in hybrid species have, however, rarely been made.

To address the relative importance of genomic contingencies and ecology for hybrid phenotypes, we investigated how diet, climate and genomic composition affect beak shape and size in a hybrid species. Our study species, the Italian sparrow, is a homoploid hybrid resulting from the interbreeding between the Spanish sparrow (*Passer hispaniolensis*) and the house sparrow (*Passer domesticus*) (Hermansen et al. 2011; ELGVIN et al. 2011; Trier et al. 2014). To be able to address the effect of genomic background, we use three island populations of Italian sparrow from each of the islands Crete, Corsica and Sicily that show strong differences in genomic composition and appear to represent independent hybridization events (Runemark et al. n.d.). In the absence of contingencies, populations experiencing the same selection pressures are expected to develop similar phenotypes (Ravinet et al. 2012; Runemark et al. 2014; 2015). Therefore, if there is strong ecological selection on the beak we expect beak size and shape to correlate with diet or climate measures despite individuals having different genomic compositions as long as contingencies are not important in the system. On the other hand, if contingencies are important we expect that island origin (reflecting genomic composition) better explains beak morphology. Diet (Grant and Grant 1996; Neto et al. 2016) and climate (Eroukhmanoff et al. 2013a; Gardner et al. 2016) have previously been found to affect beak morphology, but these factors have not been studied in genomically divergent populations. Investigating these factors jointly will shed light on whether population differences within hybrid species can be adaptive or may be restricted to values along the axis of parental divergence. We used stable isotopes as a proxy for diet, a set of climatic variables previously shown to influence beak size in the Italian sparrow (Eroukhmanoff et al. 2013a), and whole genome estimates of relative parental proportions from an earlier study on the island populations (Runemark et al. n.d.) to address which factors shape phenotypic variation in a hybrid species.

## Materials and Methods

We sampled three populations of Italian sparrows from each island of Crete, Corsica and Sicily during the spring 2013 (Fig. 1a). We caught 10-38 birds in each population (see Supplementary Table 1 for sample sizes and sex) using mist netting, and took digital pictures of the right side of each birds’ head with a Nikon D-500 16.2 megapixel camera. The background was millimeter squared paper, and we ensured that the head of the bird was not tilted. Geometric morphometrics was used to analyze beak shape. We used the thin-plate spline based programs developed by (Rohlf 1998) for file conversion (tpsUTIL) and digitization of landmarks (tpsDIG2). Five homologous landmarks were placed on the beak, and we drew an outline with 7 equidistant points, i.e. semi-landmarks to further capture beak shape (Supplementary Figure 1). PAST (Hammer et al. 2001) was used to estimate Relative Warps (RWs) and centroid size. Relative warps are principal components of shape (Zelditch et al. 2004), and were extracted (n=32) and imported to R for further analysis. All further statistical analyses were performed in R (team n.d.). As feathers for female stable isotope analysis were only sampled for one population on each island, we performed all tests on two additional datasets to ensure that this did not bias our findings. The two data sets included only set with one population from each island where both males and females were sampled, and set with all nine populations where only males were sampled.

**Figure 1.**
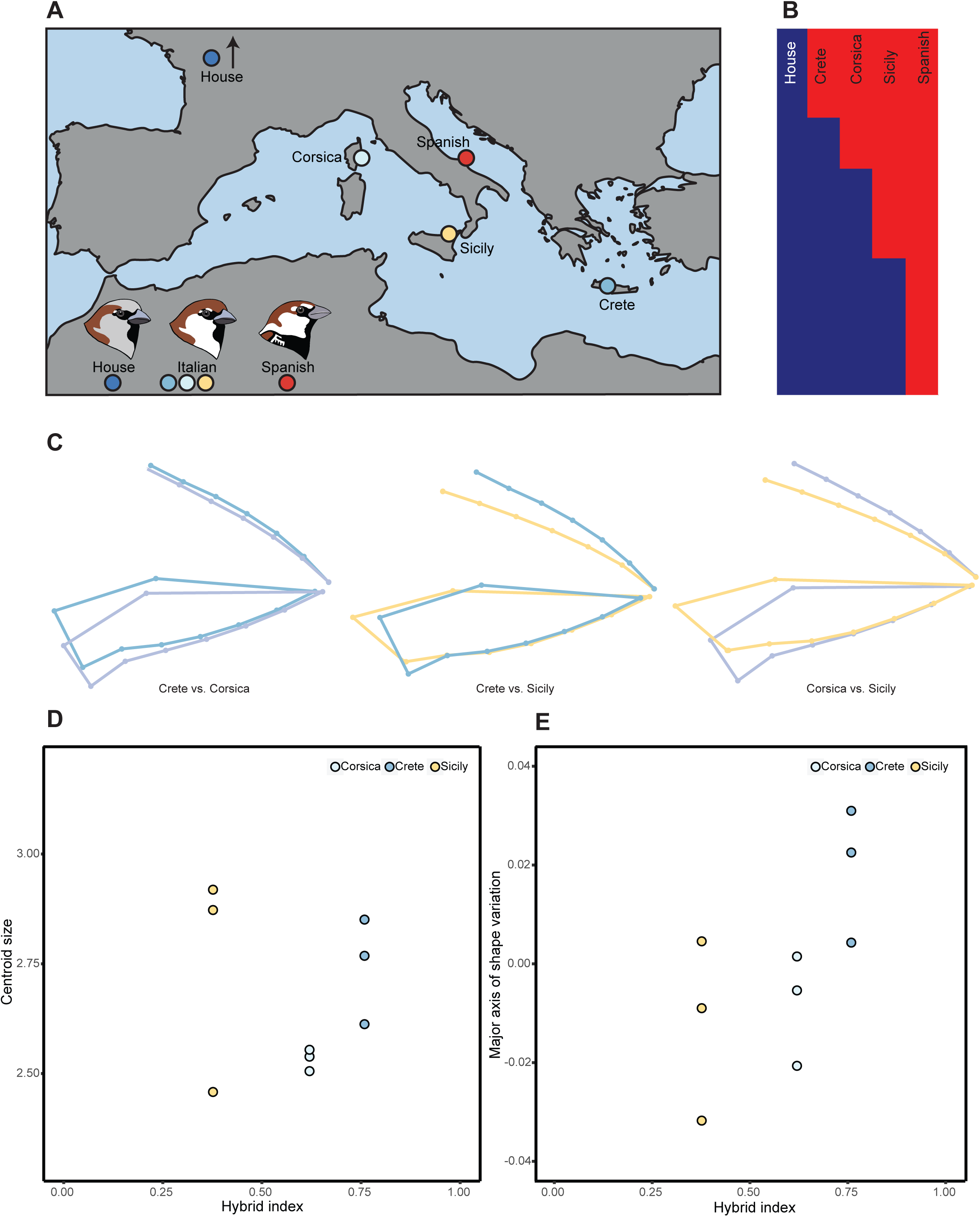
Description of the study system and beak morphology divergence. The Italian sparrow is a hybrid between the house sparrow and the Spanish sparrow. **A)** Independent, genetically divergent populations are found on the islands of Crete, Corsica and Sicily (Runemark et al. n.d.). Three populations were sampled from each island, see Supplementary Table 1 for coordinates. **B)** Hybrid index, e.g. the probability of house sparrow origin based on whole genome data, differs between populations with Crete being most house-like and Sicily most Spanish-like. **C)** Pair-wise mean beak shape differences between populations (size differences are scaled). **D)** Population divergence in size is not merely reflecting island of origin. **E)** The major axis of shape variation is not predicted by island of origin either.

**Table 1.** Model selection table. Dependent variable, replicated unit (reflecting whether the analysis was performed at the population or individual level), explanatory factors included in the model, whether a random factor was included and the AICc values and, when relevant, importance values the model selection was based on are included. Each set of tests has its own headline, and the best model is presented in bold text.

| Dependent variable(s) | Replicated unit | Factor | Random Factor | AICc | Importance AICc |
| --- | --- | --- | --- | --- | --- |
| **Size analyses at the population level** |  |  |  |  |  |
| Size | Population | $\delta^{13}\text{C}$ | None | 3.63 | 0.085 |
| Size | Population | $\delta^{15}\text{N}$ | None | 3.1 | 0.129 |
| Size | Population | Annual_Temp | None | 2.3 | 0.18 |
| Size | Population | Annual_Prec | None | 2.7 | 0.155 |
| **Size** | **Population** | **Temp_Seas** | **None** | **-4** | **0.84** |
| Size | Population | Prec_Seas | None | 3 | 0.135 |
| Size | Population | Island | None | 7.2 | 0.019 |
| **Size analyses at the individual level** |  |  |  |  |  |
| Size | Individuals | $\delta^{13}\text{C}$ | Population | -140.6 | <0.01 |
| Size | Individuals | $\delta^{15}\text{N}$ | Population | -142.5 | 0.0126 |
| Size | **Individuals** | **$\delta^{15}\text{N}+\text{HI}+ \delta^{15}\text{N}\times\text{HI}$** | **Population** | **-166.58** |  |
| Size | Individuals | $\delta^{15}\text{N}+\text{Sex}+\text{HI}+$<br>$\delta^{15}\text{N}\times\text{HI}+ \text{HI}\times\text{Sex}$ | Population | -165.59 | |
| Size | Individuals | $\delta^{15}\text{N}+\text{HI}+\text{Sex}+\text{HI}\times\text{Sex}$ | Population | -164.61 | |
| **Univariate shape analyses at the population level** |  |  |  |  |  |
| **RW1** | **Population** | **$\delta^{13}\text{C}$** | **None** | **-42.4** | **0.737** |
| RW1 | Population | $\delta^{15}\text{N}$ | None | -36 | 0.103 |
| RW1 | Population | Annual_Temp | None | -41.8 | 0.678 |
| RW1 | Population | Annual_Prec | None | -36.1 | 0.106 |
| RW1 | Population | Temp_Seas | None | -35.7 | 0.088 |
| RW1 | Population | Prec_Seas | None | -40.2 | 0.476 |
| RW1 | Population | Island | None | -36 | 0.103 |
| **Univariate shape analyses at the individual level** |  |  |  |  |  |
| **RW1** | **Individuals** | **$\delta^{13}\text{C}$** | **Population** | **-337.04** | **0.176** |
| RW1 | Individuals | $\delta^{15}\text{N}$ | Population | -311.9 | <0.01 |
| RW1 | Individuals | $\delta^{13}\text{C}+\text{HI}$ | Population | -335.81 | |
| RW1 | Individuals | $\delta^{13}\text{C}+\text{Sex}$ | Population | -334.94 | |
| **Multivariate shape analyses at the population level** |  |  |  |  |  |
| RW1-4 | Population | $\delta^{13}\text{C}$ | None | -240.09 | |
| RW1-4 | Population | $\delta^{15}\text{N}$ | None | -231.09 | |
| RW1-4 | Population | Annual_Temp | None | -239.47 |  |
| RW1-4 | Population | Annual_Prec | None | -243.55 |  |
| RW1-4 | Population | Temp_Seas | None | -239.42 |  |
| RW1-4 | Population | Prec_Seas | None | -237.15 |  |
| **RW1-4** | **Population** | **Annual_Prec$\times\text{HI}$** | **None** | **-312.43** |  |
| RW1-4 | Population | Annual_Prec+HI | None | -284.05 |  |
| **Multivariate shape analyses at the individual level** |  |  |  |  |  |
| RW1-4 | Individuals | $\delta^{13}\text{C}$ | Population | 5792.000 | |
| **RW1-4** | **Individuals** | **$\delta^{15}\text{N}$** | **Population** | **5788.888** |  |
| RW1-4 | Individuals | $\delta^{15}\text{N}+ \text{HI}$ | Population | 5790.069 | |
| RW1-4 | Individuals | $\delta^{15}\text{N}+ \delta^{13}\text{C}$ | Population | 5788.191 | |
| RW1-4 | Individuals | $\delta^{15}\text{N}\times\text{HI}$ | Population | 5792.283 | |
| RW1-4 | Individuals | $\delta^{15}\text{N}\times\delta^{13}\text{C}$ | Population | 5789.635 | |
| RW1-4 | Individuals | $\delta^{15}\text{N}\times\delta^{13}\text{C}\times\text{HI}$ | Population | 5792.25 | |
| RW1-4 | Individuals | $\delta^{15}\text{N}+ \delta^{13}\text{C}+ \text{HI}$ | Population | 5790.186 | |
| RW1-4 | Individuals | $\delta^{15}\text{N}\times\delta^{13}\text{C}+ \text{HI}$ | Population | 5790.897 | |

First, we established whether there were significant differences in beak size and shape using centroid size and the four RWs deviating from the noise floor (Supplementary Table 2) as response variables in ANOVA and MANOVA, respectively. We tested both for the presence of overall population variation and for variation among populations within islands using models with population nested within island.

Next, we investigated which factors best explain size and shape variation. We used stable isotopes as a proxy for dietary differentiation. The combination of δ^15^N and δ^13^C isotope ratios provide a comprehensive picture of diet; δ^15^N differentiation increases with each trophic level and is indicative of the trophic position in the food web (reviewed in (Caut et al. 2009). δ^13^C varies between C^3^ and C^4^-plants (Fry 2006) and δ^13^C ratios in plants decrease with rainfall (Stewart et al. 1995; Ferrio and Voltas 2005); therefore δ^13^C values are a proxy for dietary source. To obtain stable isotope values, we sent great covert feathers sampled during spring (March-June; 1 mg +/- 0.2 mg finely cut samples in tin capsules, article no. D1008, Elemental Microanalysis, Devon, UK) for δ^13^C and δ^15^N analysis at UC Davis Stable Isotope Facility. As the aim was to examine population differences, and since sparrows feed on a wide variety of resources, we did not attempt to examine isotopic contents of potential diet items, but rather whether diet differed. As baseline climatic differences could affect isotopic contents, we examined whether values clustered within islands. This was not the case (data not shown), and dietary differences were therefore not overrun by baseline signatures. We also used climatic factors previously shown to correlate with a beak size measurements in Italian sparrows (Eroukhmanoff et al. 2013a) as proxies for local climate. We extracted climate variables; annual temperature, annual precipitation, temperature seasonality and precipitation seasonality from the Worldclim database (Hijmans et al. 2005) using the R-packages raster (Hijmans and van Etten 2016), rgdal (Bivand et al. 2016) and foreach (Calaway et al. 2015). Population hybrid index estimates were retrieved from (Runemark et al. n.d.). They were based on a whole genome ADMIXTURE analysis (Alexander et al. 2009), and the mean population probability of house sparrow ancestry was used as an index. The genomic hybrid index differs between all islands, and if beak shape similarity to the parent species corresponds to genomic resemblance, this would be an important factor in the models. Thus, these variables were used as explanatory factors in our models.

Centroid size and shape were used as dependent variables. Two models were run for shape: One with only the main axis of divergence, RW1, explaining > 60% of the variation in shape, and another including all four relative warps that deviate from the noise floor. As climate is identical for all individuals within a population whereas diet may vary between individuals within a population, one population level dataset was created to address the effects of both diet and climate, and one individual level dataset soley with individual diet estimates. To test which models best explain size and shape we used a model selection framework based on applicable information criteria.

### Population level analyses

For the population level analyses, we first tested which ecological factors best explain population divergence in beak size and shape. For the models with centroid size as dependent variable, AICc and importance were estimated using the R-package MuMIn (Barton 2016). AICc is a version of Akaike’s Information Criterion, (Akaike 1974) which is especially suited for small datasets, and importance is the sum of Akaike weights (Wagenmakers and Farell 2004) 193 over all models including the explanatory variable. The variables with highest importance were used in subsequent models. We then tested which of all possible models best explained data based on AICc with sex, hybrid index and their interactions as explanatory variables. The same model was repeated for size, except RW1, reflecting a change from wide to a narrow basal part of the beak (Supplementary Figure 2) was used as a response variable. For the shape analysis including the four main RWs (Supplementary Figure 2), selection was based on AIC on MANOVA, first using models including only one climate or diet variable, and then testing if adding sex, hybrid index and/or their interactions improved the model.

### Individual level analyses

For the individual level dataset, model selection was performed as in the population level analyses, but on mixed models with population as a random factor with centroid size and RW1 as response variables, respectively. We used the lmer command from the R-package lme4 (Bates et al. 2016) for these analyses. We first tested which of the ecological variables best explained the model, and then explored whether adding hybrid index, sex and/or the interactions improved the model in the same manner as the population level analyses. To retrieve *F*- and *P*-values for the mixed models, we used the mixed function supplied in the R package afex (Singman et al. 2016).

For the shape analyses including all four main RWs, the R package MCMCglmm (Hadfield 2010) was used. When the number of groups is low the posterior distribution of the variance becomes increasingly tail-heavy, causing poor mixing of the MCMC chain. To mitigate this, we used parameter expansion (Hadfield 2010), on the MCMCglmm algorithm to speed up the rate of convergence in the MCMC chain. This entails using information from a run with an uninformative prior on the same data to choose proper values for the prior means and prior covariance matrix (alpha mean and variance) to be specified in the parameter expanded run. We then used a Cauchy prior as recommended for the parameter expanded run (Hadfield 2010), with the alpha variance set to the square of the standard deviation in the posterior distribution from the uninformative prior. The posterior sampling was run for 200 000 iterations with a burn-in of 40 000 and a thinning of 100. The MCMC-chain was plotted and inspected for proper mixing, and autocorrelation remained low (< 0.1) between successive samples in the chain. Three chains were run to ensure consistency in parameter estimation. Model selection for these models was performed based on DIC.

Finally, we addressed whether the variation among Italian sparrow populations is aligned with the axis of parental divergence, or if the phenotypic values attained deviate from this. We used PAST (Hammer et al. 2001) to estimate RWs and centroid size for a dataset including both the Italian sparrow populations and one reference population of each parent species. For size, we used an ANOVA with centroid size from this analysis as response variable, and species as a grouping factor. For shape, we performed a discriminant function analysis based on parental values only in PAST (Hammer et al. 2001), and then transformed RW scores for the Italian sparrow individuals into discriminant scores using the factor loadings of the discriminant axis between parent species. We then tested whether the position along the score axis was affected by hybrid index, thus reflecting a correlation between genomic and phenotypic similarity to the parent species using a linear regression. This will shed light on whether genomic composition constrains phenotypic adaptation within the Italian sparrow.

## Results

Sex did not significantly affect beak size or shape, and was not included in any of the best models for the dataset with both females and males in all populations (Supplementary Table 3), therefore we proceeded with our analyses using the full dataset.

### Population divergence in hybrid index, beak size and beak shape

Independent island populations from Crete, Corsica and Sicily differ in the proportion of the genome inherited from house sparrow (Runemark et al. n.d.)(Figure 1b). Beak size varies between populations (size: F_8,127_=18.75; *P*=2e-16; shape: F_32,508_=2.81; *P*=1.05e-06; (Fig. 1c-e). These differences persist if population is nested within island both for size (island: F_2,127_=22.56, *P*=4.12e-09; population nested within island F_6,127_=17.48, *P*=1.13e-14) and shape (island:F_8,250_=6.94, *P*=2.97e-08; population nested within island: F_24,508_=1.69, *P*=0.022). The presence of significant variation within islands shows that differences do not merely reflect genomic composition (Fig. 1c-e), but are influenced by other factors.

### Beak size

Temperature seasonality was the factor best explaining population divergence in beak size, and had ΔAICc of more than 6 to the second best model (Tables 1-2; Figure 2a). As all individuals in a population experience the same climate, we also tested which factors affect beak size at the individual level, excluding climate variables. The best model for individual variation includes δ^15^N, genomic hybrid index and the interaction between these factors (Figure 2b), reflecting that δ^15^N changes do not affect individual beak size in the same manner across islands. Two models were within ΔAICc of 2 of this best model (Tables 1-2). One included sex and the interaction between sex and hybrid index in addition to the abovementioned factors, whereas the other included δ^15^N, genomic hybrid index, sex and the interaction between genomic hybrid index and sex. Hence, patterns of individual beak size variation are complex and no clear best explanatory variables emerge.

**Figure 2.**
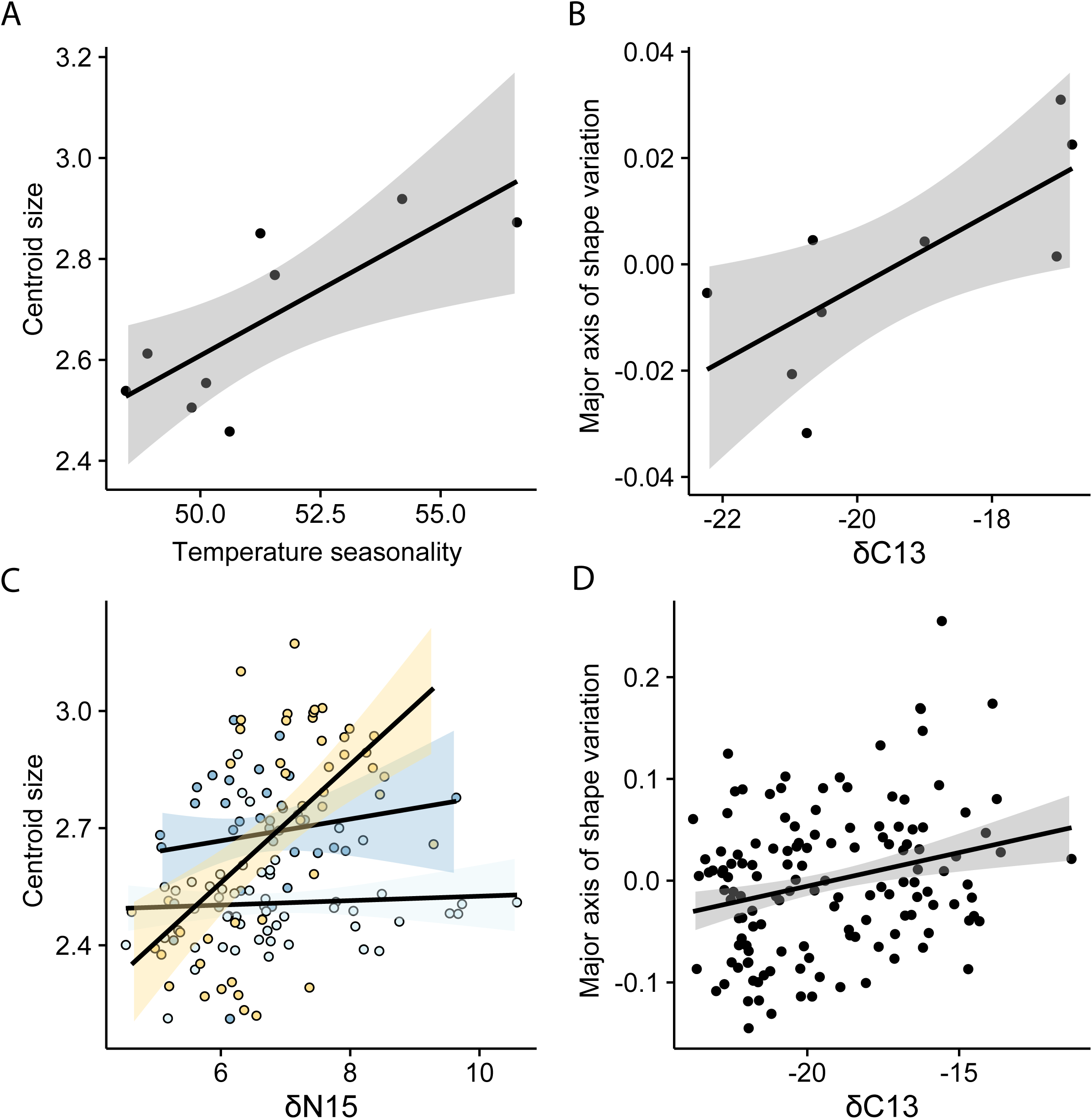
Factors best explaining size and shape variation. **A)** Temperature seasonality is the best predictor of centroid size at the population level, and the relationship is highly significant (*F*_1,7_=10.26; *P*=0.015; R^2^=0.59). **B)** δ^13^C best explained population divergence along the main axis of variation (*F*_1,7_=8.01; *P*=0.025; R^2^=0.47). **C)** At the individual level, centroid size was best explained by a model including both δ^15^N and genomic hybrid index and their interaction, as the relationship between δ^15^N and centroid size varied between islands (model R^2^=0.72). **D)** Individual level shape divergence along the axis of largest variation was, as for the population level, best explained by δ^13^C (R^2^=0.07).

**Table 2.** Properties of the best models. *F*-values, degrees of freedom, *P*-values (for lm and lmer models) and pmcmc values (for the MCMCglmm model) and model R^2^ for the models where it is applicable.

| Dependent variable | Factor | Estimate | *F* | DF, error DF | *P*/pmcmc | Model $R^2$ |
| --- | --- | --- | --- | --- | --- | --- |
| **Population level analyses** |  |  |  |  |  |  |
| **Size** | Temperature seasonality | 0.052 | 10.26 | 1, 7 | 0.015 | 0.5945 |
| **Warp1** | $\delta^{13}\text{C}$ | | 8.01 | 1, 7 | 0.025 | 0.467 |
| **Warp1-4** | Annual precipitation | -0.002 | 2.78 | 5, 3 | 0.21 | 0.8225 |
|  | HI | 0.025 |  |  |  |  |
|  | Annual precipitation×HI | -4.56e-05 |  |  |  |  |
| **Individual level analyses** |  |  |  |  |  |  |
| **Size** | $\delta^{15}\text{N}$ | -0.092 | 2.94 | 1, 128.89 | 0.09 | 0.7209 |
|  | HI | -1.28 | 2.72 | 1, 43.75 | 0.11 |  |
| | $\delta^{15}\text{N} \times \text{HI}$ | 0.16 | 2.89 | 1, 128.95 | 0.09 | |
| **Warp1** | $\delta^{13}\text{C}$ | | 4477<br>4.00 | 1, 29 | 0.008 | 0.0712 |
| **Warp1-4** | $\delta^{15}\text{N}$ | 8.41 | NA | 1, 129 | 0.0368 | NA |

### Beak shape: the major axis of divergence

The best model for population divergence along the main axis of shape variation, reflecting a change from a wide to a narrow basal part of the beak (Supplementary Figure 2), included only δ^13^C, and explained the data significantly better than the second best model (ΔAICc > 2; Tables 1-2; Figure 2c). Individual level variation in beak shape was also best explained by δ13C differences (Figure 2d), with ΔAICc to the second best model of >4 (Tables 1-2).

### Beak shape: all significant axes of divergence

The first four RWs reflecting beak shape variation deviated from the noise floor (Supplementary Table 1; Supplementary Figure 2). The model best explaining this shape variation included annual precipitation, genomic hybrid index and the interaction between these terms (Supplementary Figure 3a-d; Tables 1-2). We also tested which factors affect beak shape at the individual level, excluding climate variables. Individual shape differences were best explained by a model including only δ^15^N (Supplementary Figure 3e-h; Tables 1-2).

### Parental phenotypes and the extent of genomic contingencies

We estimated the axis discriminating the parent species based on the four RWs deviating from the noise floor (Supplementary Table 4), and scored the hybrids on this axis. We found a significant correlation between hybrid index and score along the parental axis of variation (estimate=9.08±2.95; *F*_1,199_ =9.50; *P*=0.002; R^2^=0.05), implying that populations that are genomically similar to house sparrows also have a more house sparrow like beak shape. Breaking up shape into the individual axes of variation, we find intermediacy and hence potential constraints only in the third and fourth shape component, while Italian sparrows attain values outside of the parental range for the first and second (Supplementary Figure 3). Centroid size was nearly significantly correlated with hybrid index (estimate=0.11±0.058; *F*_1,199_ =3.64; *P*=0.058; R^2^=0.01; Supplementary Figure 4).

## Discussion

Both beak size and beak shape vary significantly between Italian sparrow populations, as well as between islands. Interestingly size and shape are not best explained by the same factors at the population level. While beak size is strongly affected by temperature seasonality, the main axis of beak shape variation is best explained by variation in carbon isotopic ratios. Although ecological factors best explain beak shape along the major axis of variation, beak shape divergence for all significant axes of variation is significantly affected by genomic hybrid index, reflecting island of origin and potentially contingencies. The fact that there is a correlation between position along the discriminant axis separating the parent species’ shape and the genomic similarity to the parent species is also consistent with a role for contingencies. Patterns of individual axes of variation do, however, suggest that there may be contingencies in some, but not all, directions of variation.

There are various reasons temperature regime could affect beak size. Temperature variation could affect the size spectrum of the available diet. There is mounting evidence that beaks play an important role in thermoregulation, as blood flow through the network of supportive blood vessels beneath the keratinized surface is augmented at high temperatures and restricted in the cold (Symonds and Tattersall 2010; Campbell-Tennant et al. 2015). For instance, beak sizes vary as expected from Allen’s rule (Allen 1877), which posits that the relative size of body extremities is smaller in colder environments, for ectotherms to reduce thermoregulatory costs (Symonds and Tattersall 2010). Even if the effect of smaller beaks cannot explain a high proportion of total heat loss, as in the toucan (Tattersall et al. 2009), using the beak for thermoregulation could potentially be important during summers on these arid Mediterranean islands. Furthermore, the fitness advantage of large bill size could differ depending on local temperature profiles and humidity, even in small passerine birds (Gardner et al. 2016). Individual level divergence is affected by a more complex combination of factors, and no clear best model emerged, although both nitrogen isotopic composition and genomic hybrid index were included in all models. This relationship could therefore be complex and involve many factors of small effect or variables that we have not measured.

Annual precipitation pattern is the ecological factor best explaining beak shape. Interestingly, both general beak shape as well as how precipitation patterns affect beak shape, are significantly affected by genomic hybrid index. Precipitation patterns could affect seed size (Moles et al. 2005) and the hardness of seeds (Mohamed-Yasseen et al. 1994). Seed size is known to affect beak size evolution in passerines (Grant and Grant 1993), including in sparrows (Riyahi et al. 2013). In addition, beak shape affects bite force (Herrel et al. 2005), and the correlation between annual precipitation and beak shape could reflect adaptation to deal with harder seeds. As the significant genomic hybrid index term and interaction between genomic hybrid index and annual precipitation suggest, the significant genomic hybrid index term may reflect a genomic contingency in form of an island specific beak shape and beak shape variation structure implying responses to the same selective environment differ,. The same increase in annual precipitation does hence not result in the same shape response across the islands. The correlation between genomic similarity to a parent species and shape similarity to that species suggests that this potentially could be due to genomic contingencies.

Nitrogen isotopic composition is the only factor in the model that best explained individual beak shape differences along all dimensions. Differentiation in isotopic composition between a consumer and dietary items is low, predictable and conserved across trophic levels (i.e. typically 1 ‰ difference) (Peterson and Fry 1987). Therefore it allows for accurate discrimination of dietary contributions from different nitrogen sources (Newsome et al. 2007). Thus stable isotope signatures may reflect dietary differences in birds, which in turn may also influence beak shape (Neto et al. 2016). Beak specialization for foraging in different selective regimes are well-established in birds (Grant and Grant 1996; Benkman 2002; 2016). The Italian sparrow is an opportunistic human commensal species, which feeds both on wild seeds, crop plants and insects. Specialization enabling foraging on prey from different trophic levels, or differences in proportions in individual diet within populations could potentially explain the effect of nitrogen isotopic composition on beak shape. Although all sampled individuals were breeding adults, stable isotope composition reflects diet at molt the previous autumn, and the birds could have belonged to different age classes at this point in time.

Interestingly, one of the genes that was most divergent between Crete and Sicily in a study of the genomic composition of the island populations was FGF10 (Runemark et al. n.d.), a candidate gene for beak shape shown to be important in beak divergence in Darwin’s Finches (Lamichhaney et al. 2015). Together with the ecological differences and correlated beak morphology divergence, this genomic signature of selection on the genes affecting the phenotype makes a strong case that the sorting of parental variants allows hybrid species to locally adapt.

The island populations of Italian sparrow from this study have contingencies in the proportion of inheritance from each parent species (Runemark et al. n.d.), resulting from mosaic patterns of parental inheritance or conditions during initial hybridization and genome stabilization c.f. (Eroukhmanoff et al. 2013b). We find that hybrid taxa are intermediate between parent species for both size and shape, although not for all shape components. This is consistent with the pattern predicted for traits where directional selection contributes to parent species differences in which hybrids are expected to differentiate along the parental axis of divergence (Bailey et al. 2013). Furthermore, the proportion of the parental genome inherited from each species, here measured as hybrid index, was significantly correlated with similarity to the parent species beak shape. Taken together, that genomic hybrid index is involved in the best model explaining population divergence in beak shape and is significantly correlated with position along the parental axis of variation suggests that constraints may affect evolutionary trajectories and evolutionary potential following hybridization. There are, however, two shape dimensions that are transgressive. This demonstrates a release of parental constraint for some components of shape, and is consistent with the predicted patterns of divergence for traits under stabilizing selection in the parents (Bailey et al. 2013).

In conclusion, this study provides evidence of adaptive local divergence within a hybrid species, but shows that genomic contingencies could affect the evolutionary potential to respond to selection in a hybrid species. Size and shape divergence are best explained by different selective factors, with temperature patterns affecting size and precipitation patterns and proportion inherited from different parent species predicting shape. Interestingly, we only find evidence for constraint in shape and not in size, consistent with patterns in the fossil record suggesting that size is more evolvable than shape (Hunt 2007).

